# Inference of Transmission Network Structure from HIV Phylogenetic Trees

**DOI:** 10.1101/059865

**Authors:** Federica Giardina, Ethan Obie Romero-Severson, Jan Albert, Tom Britton, Thomas Leitner

## Abstract

Phylogenetic inference is an attractive mean to reconstruct transmission histories and epidemics. As the interest lies in how HIV-1 spread in a human population, many previous studies have ignored details about the evolutionary process of the pathogen. Because phylogenetics investigates the evolutionary history of the pathogen rather than the spread between hosts per se, we first investigated the effects of including a within-host evolutionary model in epidemiological simulations. In particular, we investigated if the resulting phvlogenv could recover different types of contact networks. To further improve realism, we also introduced patient-specific differences in infectivitv across disease stages, and on the epidemic level we considered incomplete sampling and the age of the epidemic. Second, we implemented an inference method based on approximate Bayesian computation (ABC) to discriminate among three well-studied network models and jointly estimate both network parameters and key epidemiological quantities such as the infection rate. Our ABC framework used both topological and distance-based tree statistics for comparison between simulated and observed trees. Overall, our simulations showed that a virus time-scaled phvlogenv (genealogy) may be substantially different from the between-host transmission tree. This has important implications for the interpretation of what a phvlogenv reveals about the underlying epidemic contact network. In particular, we found that while the within-host evolutionary process obscures the transmission tree, the diversification process and infectivitv dynamics also add discriminatory power to differentiate between different types of contact networks. We also found that the possibility to differentiate contact networks depends on how far an epidemic has progressed, where distance-based tree statistics have more power early in an epidemic. Finally, we applied our ABC inference on two different outbreaks from the Swedish HIV-1 epidemic.

## Introduction

Infectious diseases that are directly transmitted spread over contact networks; each individual or host can be represented by a node with a finite set of contacts/edges to whom they can pass the infection. The structure of these networks is a major determinant of the pathogen transmission dynamics and possible control strategies [1], For example, it has been suggested that human sexual contact networks are characterized by a power-law degree distribution [2] which, in a specific range of the scaling exponent, results in an infinite variance of the network’s degree distribution. This implies the absence of an epidemic threshold which makes prophylactic strategies for sexual transmitted diseases very challenging.

The main issue in contact network epidemiology has been the difficulty of collecting individual and population-level data needed to develop an accurate representation of the underlying host population’s contact structure. This has led to an interest in methods to infer information about host contact networks from epidemic data. Previously, Britton and O’Neill [3] estimated the parameters of an Erdõs-Rénvi network and a stochastic epidemic process on it using epidemic data consisting of recovery times of infected hosts and Groendvke et al, extended the approach to exponential-family random graph models [4] using covariate information [5], The use of other common epidemiological measures such as the basic reproduction number (*R*_0_), epidemic peak size, duration and final size, has been shown to be effective in classifying the degree of heterogeneity in a population’s unobserved contact structure [6].

During the course of a given epidemic, the disease spreads over a subset of edges in the social network forming a subgraph that is the realized transmission history. Keeping track of who transmits to whom and assuming that every individual may be infected only once and by only one other individual, such a transmission history can be represented as a rooted tree (*transmission tree*) [7], However, full transmission histories are rarely observed and commonly available epidemiological data such as infected people diagnosis-reeoverv times may provide information on who was infected, when, and for how long, but it cannot provide information on who acquired infection from whom.

Since pathogens evolve over a transmission history, the analysis of pathogen genetic sequences taken from different hosts provides a way to infer the most likely donor and recipient [8] introducing constraints on the space of possible transmission trees, which are a trace of the underlying contact network, Phylodvnamies [9] focuses on linking methods of phylogenetic analysis with epidemiological models under the assumption that if the evolution of a pathogen occurs sufficiently fast, transmission histories become recorded in the phvlogenv of the pathogen population (*phylogenetic tree*).

Phylodvnamie analyses on HIV have shown that asymmetry in viral phytogenies may be indicative of heterogeneity in transmission [10], Networks with more heterogeneous degree distributions yield transmission trees with smaller mean cluster sizes, shorter mean branch lengths, and somewhat higher tree imbalance than networks with relatively homogeneous degree distributions, However, it has been argued that these direct effects are relatively modest for dynamic networks [11] or if only a small fraction of infected individuals is sampled [12], Also, factors other than contact rate, such as high infectiousness during acute infection, may have a more dramatic impact on asymmetry [12].

However, previous studies as well as more recent papers [13–15], all assume that the HIV unobserved transmission tree is identical to the reconstructed time-scaled phvlogenv (*virus genealogy*), i.e. the internal nodes of the genealogy correspond to transmission events between hosts over time and within-host diversity is fundamentally ignorable. This is unrealistic since all the coalescent events in a pathogen phvlogenv occur within hosts, pushing the genealogy node heights further back in time than the nodes of the transmission tree [16], In addition, the order of coalescent events may not correspond to the order of transmission events but reflect instead within-host dynamics [17].

The objective of this study was to include within-host evolution, disease stage and individual specific transmission rates to improve the realism of social network reconstruction. We simulated epidemic spread on three network types and investigated the behavior of several tree statistics, including both topological imbalance measures and tree-based distance measures. In addition, we investigated the effect of varying epidemic size, varying sampling proportion as well as heterochronous sampling on the tree statistics. Finally, we analyzed data from two different epidemiological sets of spread among injecting drug users (IDU) in the Swedish HIV epidemic using approximate Bayesian computation (ABC) for network model choice and parameter inference following the algorithm defined in [18], We found that virus geneaolo-gies can differ from the underlying transmission tree in both topology and branch length and, therefore, meaningful inference of social networks needs to include within-host evolution.

## Materials and Methods

### Simulation of transmission history

**Networks** We considered three different network models to represent population structure: the Erdos-Renyi (ER) random graph [19], the Barabasi-Albert (BA) graph [20] and the Watts-Strogatz (WS) graph [21] with low rewiring probability (Figure 1). These three networks are characterized by different degree distributions and amount of clustering. The degree of a node in a network is the number of connections to other nodes it has and the degree distribution is the probability distribution of these degrees over the whole network.

The ER model generates networks with Poisson degree distributions, i.e. 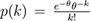 The BA model is generated by using a linear preferential attachment algorithm that produces scale-free networks with a power-law degree distribution *p*(*k*) ∝ *k^−α^* with *α* = 3. The WS model has a Dirac degree distribution centered at K (all nodes have the same degree) when the rewiring probability tends to 0. If the rewiring probability tends to 1, the degree distribution is Poisson. For intermediate values, the shape of the degree distribution has a pronounced peak at *k* = *K* and decays exponentially for large |*k − K* |. A WS network is characterized by a relatively homogeneous structure, as all nodes have more or less the same degree, and by a high degree of local clustering as opposed to ER and BA networks.

**Figure 1:**
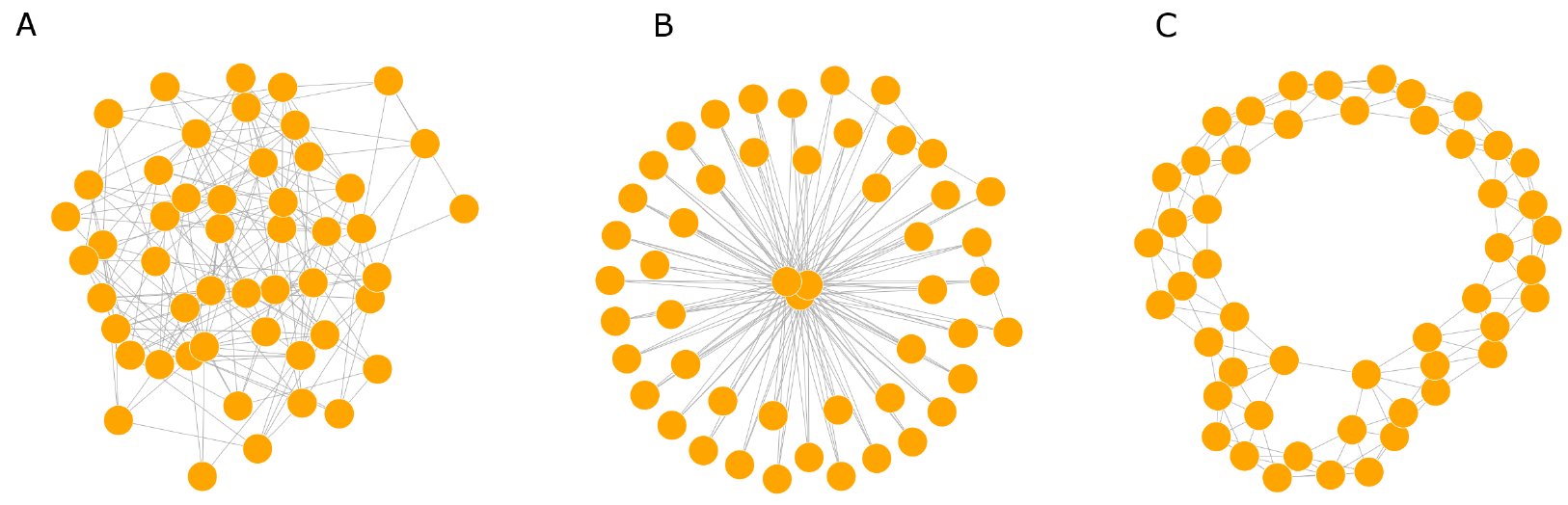
Prototypic network structures. A. Erdõs-Rénvi network network (ER), B. Barabási-Albert network (BA), C. Watts-Strogatz network (WS). To illustrate the typical configurations, all networks are generated with the same mean degree (6) in this example.

**Epidemic model** We simulated outbreaks from a susceptible-infected-removed (SIR) type dynamic [22] of HIV spread in the susceptible population of each social network. We compared differences among the transmission trees obtained by simulated epidemic spread under four increasingly more realistic transmission (Figure S1) and evolutionary model specifications (Figure S2).

The first model specification assumed that the rate of transmission per contact between a susceptible and an infected individual, *λ*, is constant over time. The removal rate of infected individuals *γ* is also constant over time and include both diagnosis or death events. We denoted with *p* the probability of being sampled at the moment of diagnosis (DNA sequences obtained from the virus of the diagnosed patient). We assumed that:

1. diagnosis coincides with treatment start,
2. the rate of transmission after treatment start is negligible, and
3. nobody goes off treatment.

We believe these assumptions to be reasonable for our analysis since Sweden has already achieved the 90-90-90 target set by UNAIDS in 2014 [23] according to which:

1. 90% of all people leaving with HIV will know their HIV status,
2. 90% of all people with diagnosed HIV infection will receive sustained antiretroviral therapy, and
3. 90% of all people receiving antiretroviral therapy will have viral suppression [24],

In the second model specification we considered three stages of HIV infection (acute, chronic, and pre-AIDS), The transmission rates are dependent on the disease stage of the infected individual and denoted with *λ*_1_, *λ*_2_, and *λ*_3_ (Figure SI), We assumed the removal rate to be independent on the disease stage. The acute stage was assumed to last for a constant period of 30 days for each individual [25], the chronic stage had variable length described bv an exponential random variable T_2_ with a mean of 8 years [26] and the pre-AIDS stage lasted until death or diagnosis. The three transmission rates were calculated to preserve the individual total infectivity during their infectious period in order to make results comparable with the first model specification. This derivation is shown in Text S1.

In the third model specification we modeled individual variability of transmission rates. We did that by multiplying the constant transmission rate *λ* (as in the first model specification) with a log-normal variable *Z_i_* for each *i* individual (node) with mean −*σ*^2^/2 and variance *σ* in order to preserve the mean of *λ* (i.e. 𝔼(*λ_i_*) = *λ* since 𝔼(*Z_i_*) = 1).

The fourth model specification combines stage-specific infeetivitv with individual heterogeneity. The sampling process is modeled explicitly in each model specification.

We used Gillespie’s next-reaction method [27, 28] to simulate disease spread according to the above outlined model specifications until there are no more infectives or until a predefined number of samples. Keeping track of who-infects-whom, each epidemic simulation yields a transmission history.

### Within-host evolution model

Pathogens such as HIV record in their phytogenies a considerable amount of information about the transmission histories since mutations are typically accumulated faster than transmission occurs. The common assumption that the internal nodes of a phvlogenv correspond to transmission events between hosts over time is unrealistic because transmitted lineages must already exist in the donor at the time of transmission. Thus, neglecting the time difference between the common ancestor and the transmission event (i.e, the pre-transmission interval, [16]) will bias the estimated time of transmission backwards in time.

Furthermore, new infections may come from HIV variants derived from a latent reservoir (lineages can persist for long time in the host [29,30]), and the order of coalescent events may not correspond to the order of transmission events but reflect instead within-host dynamics [17].

To address these issues, we used a two-phase coalescent model including a linear growth from a single transmitted variant (transmission bottleneck) to a maximum population size followed by either stabilization or decline of the effective population size [17].

Let *N*(*t*) denote the population size at time *t* since seroconversion (expressed in days), such that

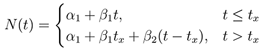

where *α_1_* is the population size (i.e. the number of virus variants in a given host) at the moment of infection, [*β_1_* is the rate of population size increase until *t_x_* (time at maximum diversity), and *β_2_* the rate of decline after the maximum. We assume *α_1_* = 1, *β_1_* = 3, *t_x_* uniformly distributed between *t_a_* = 2 and *t_b_* = 8 (years) and *β_2_* ~ *U*(*ϕ*, 0) where *ϕ* = (*N_min_ − α_1_* − *β_1_t_a_*)/(*t_M_* − *t_a_*) with *N_min_* being the minimum population size, (assumed to be 100) and *t_M_* the maximum sampling time (20 years) (Figure S2).

Virus genealogies conditional on a transmission history are simulated by generating random eoaleseenee times for each person in the tree. Random eoaleseenee times are generated from the inverse cumulative density function (derivation in [17])

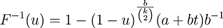

where *u* is a uniform random variate on (0,1), *t* is the current time along the forward time axis, *b* is the linear rate of change (*β_1_* or *β_2_* depending on the phase), *a* is the starting population size (1 in the first phase and *β_1_t_x_* in the second phase) and *k* is the number of extant sampled lineages in a given host. For each host we draw random values of *t_x_* and *β_2_* from the prior distributions. Starting at the last transmission or sampling event, we first move to the next event along the reverse time axis, which is either a transmission event, a phase transition, or the time at which the current host was infected. If the event is a transmission event, then *k* is incremented and a random coalescence time is generated. If that time occurs before (along the reverse time axis) the next event, then two random extant lineages in the sample are selected to coalesce; if not, then time is moved to the time of the next event. At the phase transition, *t_x_*, the parameters of the inverse cumulative density function are changed to correspond to the first phase and the process continues until the transmission time of the current host is reached. In the rare instance where more than one sampled lineage exist at the time of infection, the existing lineages are randomly coalesced with zero length branches. Finally, each individual sub-tree is jointed into a single viral genealogy according to the transmission history.

The 4 model specifications introduced in the previous section were used for simulations until the “end” of each outbreak, i.e. when there are no infectives left. We compared outbreaks of similar final size and multiple realizations of virus genealogies for each transmission history. All simulations were implemented using the statistical software E [31].

### Tree statistics

To evaluate the resulting trees from our simulations we used several tree statistics. To assess how balanced trees were, we used:

1. Sackin's Index [32], which is the average number of splits or ancestors from a tip to the root of the tree. It can be normalized according to a reference model in order to obtain a statistic that does not depend on the tree size.
2. Colless Index [33], which inspects the internal nodes, partitioning the tips that descend from them into groups of sizes *r* (on the right) and *l* (on the left of the tree), and computes the sum of absolute values |*r* −1| for all nodes. Both Sackin’s index and Colless index depend only on the topology of the tree, and they are invariant under isomorphisms and relabeling of leaves. They reach their maximum value at caterpillars (ladder-like trees), and their minimum on the maximally balanced trees, A binary tree is considered to be perfectly balanced if each internal node in the tree divides the leaves descending from it into two equally sized groups.
3. Cherries, which are defined as the number of clades with two taxa. The expected number of cherries in a tree with *n* taxa under a Yule or coalescent model is *n*/3. In an asymmetric tree (more ladder-like trees), tips tend to coalesce with branches deeper in the tree, and there are fewer cherries than expected. The number of cherries and Sackin’s index complement each other well, as the number of cherries captures asymmetry in the recent evolutionary past, while Sackin’s index captures asymmetry over the entire evolutionary history of the sample. These two measures are only weakly correlated [12].
4. The ratio between external and internal branch lengths. It has been shown that a high ratio of internal branch to external branch length occurs in ‘star-like’ trees.
5. The tree height in a virus genealogy represents the time from the first infection to the last sampling event. Since epidemics progress at different speed on different networks, heterogeneities in tree heights are expected.
6. Topological distance, obtained as twice the number of internal branches defining different bipartitions of the tips, A topological distance that takes branch lengths into account was also considered (the sum of the branch lengths that need be erased to have two similar trees).
7. The number of lineages through time normalized both in time and in number of lineages, [34].
8. The rate of branch length growth as a function of tree height (time).

We used the package R package ape (Analyses of Phylogenetics and Evolution) [35] to create and plot the phylogenies and the package apTreeshape [36] for the evaluation of some tree statistics.

### Approximate Bayesian Computation for network model selection

To further investigate how well time-scaled phylogenies can estimate the epidemic process and identify the underlying contact networks, we applied an inference framework for model selection and parameter estimation based on approximate Bayesian computation (ABC), ABC is a methodology to estimate model parameters replacing the likelihood function with a simulation-based procedure and a distance function to measure the similarity between simulated and observed data. Various ABC algorithms have been proposed, from the simple ABC-rejection [37] to ABC Markov chain Monte Carlo (MCMC) [38] and ABC based on sequential Monte Carlo (SMC) methods [18,39].

Here, we use ABC-SMC as proposed by Toni et al, [18] because it addresses some of the potential drawbacks of previous ABC algorithms, such as slow convergence rate, by sampling from a sequence of intermediate distributions, The SMC sampler introduces a number of intermediate steps decreasing iteratively the tolerance threshold e for samples acceptance. At the first iteration, *N* particles *θ*′ (representing the parameters of interest) are generated form the prior distribution and data are simulated from the model based on *θ*′ The proposed parameters are accepted if the difference between the summary statistics of the simulated data *D*′ and the observed data *D* is below the threshold *ϵ*_1_ At iteration *t* > 1, the particles are drawn from the previous population of the accepted samples at the iteration *t* − 1 (with threshold *ϵ*_t-1_) with slight perturbations. In our work, data (observed virus genealogy) and simulated trees are compared through the use of summary statistics which correspond to the above listed tree statistics.

The three network models *M* = {*WS, ER, BA*} were used to simulate outbreaks using the stage-varying infeetivitv profile with ratio 10:1 acutexhronic and patient infeetivitv variation (*σ* = 3), We assumed that network model and one network parameter were unknown. For ER, the network parameter of interest was the probability of drawing an edge between two arbitrary vertices; for BA it was the number of edges to add in each time step of the generating algorithm, and for WS it was the neighborhood within which the vertices of the lattice are connected. We also estimated the removal rate *γ* and the infection rate in the acute phase *λ*_1_ (infection rates in the chronic and immuno-compromised stage can be obtained deterministically from the acute phase infection rate). Therefore, *θ* consists of 3 parameters for each type of network and they are model specific. All remaining parameters characterizing both the network structure and the epidemic process were considered known.

The output of the algorithm were the approximations of the model *M* marginal posterior distributions *P*(*M|D*) which is the proportion of times that each model is selected in *N* samples, and the marginal posterior distributions of parameters *P*(*θ|D,M*) for the candidate models. We used a discrete uniform distribution from 1 to 3 as model prior *π*(*M*).

We chose to decrease the tolerance values following an exponential decay such that *ϵ*_t_ = *ϵ*_0_ exp(−0.5*t*) where *t* is the current sequential step, as proposed in [40]. A pilot run of 100 simulations for each model in *M* was used to define the initial thresholds. We found that converge was achieved with *T* = 10 iterations and *N* = 1000 particles per iteration. The prior distributions on the parameters *λ_1_* and *γ* were Uniform (0.0001,0.1) and (0.00025,0.1), respectively. Further details of the algorithm can be found in Text S2.

### Real epidemiological data and genealogical reconstruction

We applied the ABC inference method to the analysis of two HIV-1 sets of transmission among IDU in Sweden [41, 42], To reconstruct the time-scaled virus phylogenies from DNA sequences in both chains of transmission we used a Bayesian Skyline coalescent model in BEAST 1,8 [43], The general time reversible nucleotide substitution model was used with an uncorrelated log-normal relaxed clock and a discretised gamma distribution with four categories was used to model rate heterogeneity across the sequence. For the log-normal relaxed clock parameters, a uniform prior on the positive axis was assumed for the mean, and an exponential with mean 1/3 for the standard deviation, A Uniform prior on (0,1) was used for the nucleotide frequencies. The MCMC was run for 10 million iterations, with a 10% burn-in period and 10000 iterations. We selected the maximum credibility tree and the negative branches were set equal to zero.

## Results

### Within-host evolution affects inference of contact networks

The within-host model generates virus genealogies that are consistent with a given transmission history. An example of the within-host evolutionary impact in a small size network/epidemic is shown in Fig 2, Clearly, many virus trees are possible under any transmission history. Therefore, it is important to evaluate the additional variation within-host diversity inflicts on the epidemiological inference.

An epidemic can spread faster on EE and BA networks, thus the resulting transmission tree from a WS network includes longer times resulting in taller trees (Fig 3A), Both the unobservable transmission tree and the observable virus genealogy show the same tree height information. Other tree statistics, however, show different patterns of network discrimination based on transmission tree or virus genealogy. The proportion of cherries per taxa is slightly less informative on virus genealogies than on transmission trees (Fig 3A), In particular, while there is a decrease for EE and WS (less balanced in virus genealogies than transmission trees), it increases for BA (more balanced in virus genealogies than transmission trees), A similar pattern is seen using Sackin’s Index or Colless’ Index (EE and WS less balanced in virus genealogy, BA more balanced (Fig 3D–Figure 3E)), Overall, differences between BA and WS become more evident in virus genealogies. Because Sackin’s Index and Colless’ Index are highly correlated we will only report Sackin’s Index from now on.

The ratio of the mean internal to external branch lengths is informative about the type of network (smallest for BA, higher for EE, highest for WS), While the trends were similar in transmission trees and virus genealogies, the expected ranges overlapped for EE and WS in transmission trees, and virus genealogies showed generally smaller ratios (Fig 3C), Branch lengths increase linearly as a function of tree height during epidemic spread on both EE and BA networks. Deviations from linearity are observed for epidemic spread on WS, At the end of an epidemic, the mean branch length is constant among networks but longer in virus genealogies rather then in transmission trees (Fig 3F).

Overall, trees from EE and WS networks are more imbalanced based on virus genealogies, and because epidemic spread is much faster in a BA network the resulting virus genealogy will instead become more balanced as the virus does not have time to evolve time structure between transmission events.

**Figure 2:**
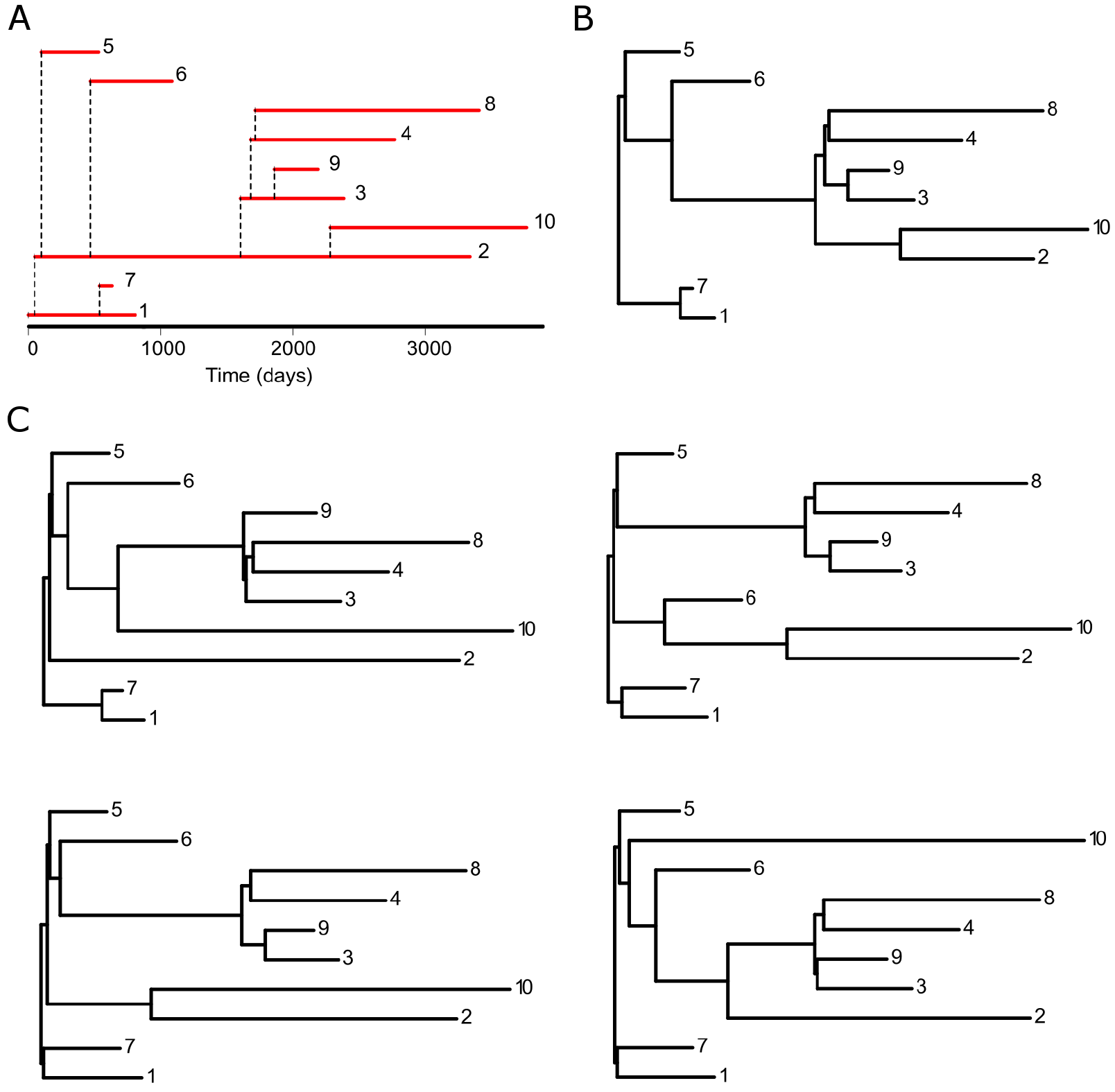
Relationship among transmission history, transmission tree and virus genealogy. For a given transmission history between hosts (A), *we* can construct a binary representation, i.e. the transmission history (B), The lower panels (C) show 4 possible virus genealogies of this transmission history invoking our within-host population model.

### Contact network heterogeneity becomes less evident under stage varying infectivity

Infectivity is known to vary across pathogenesis [44, 45], Thus, rather than assuming a constant transmission rate throughout an infected person’s disease stages, we tested 7 different infectivity profiles varying the ratio between the acute and chronic transmission rates and measured how they affected network model discrimination. The transmission rate in the pre-AIDS stage was held constant.

**Figure 3:**
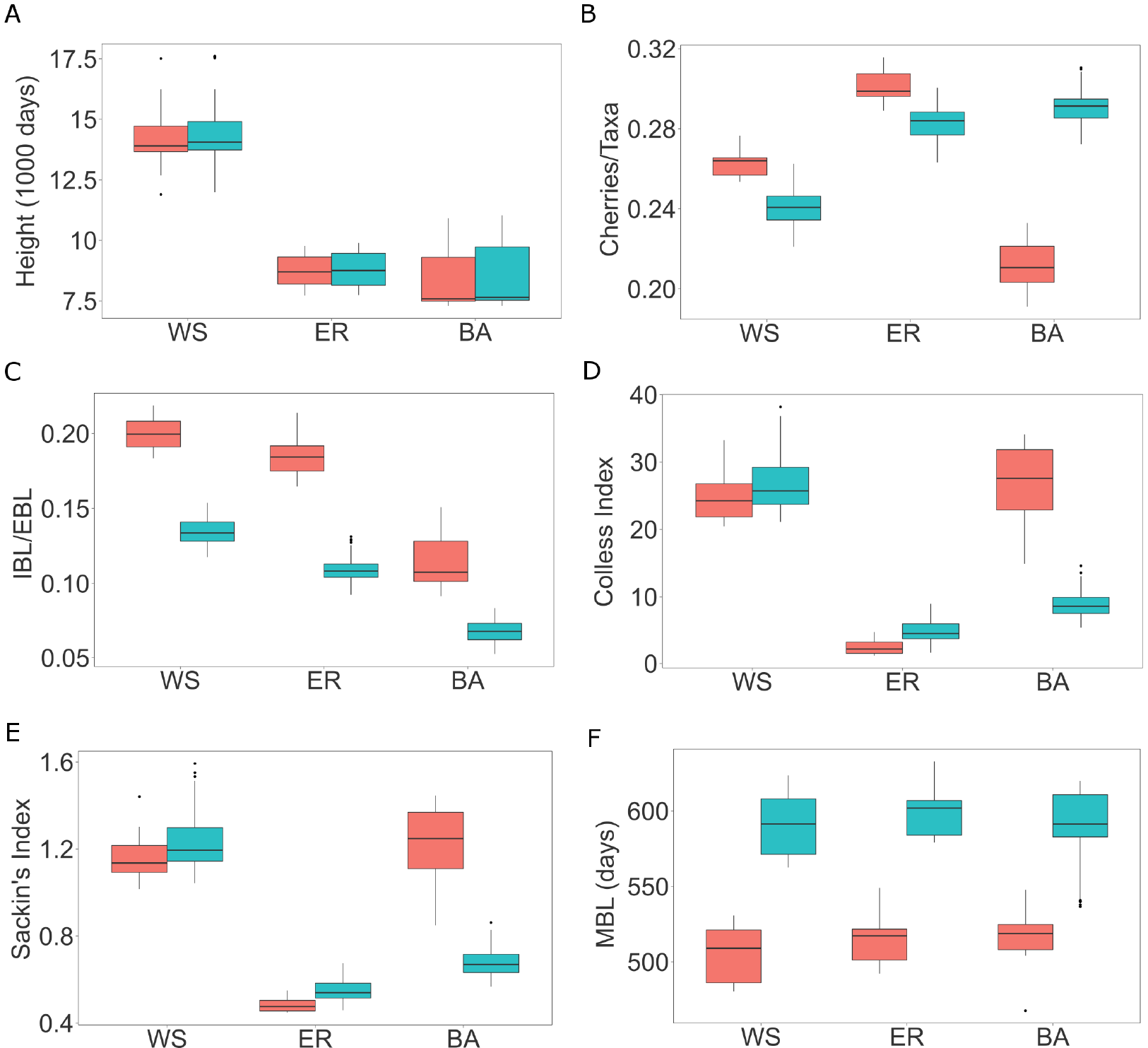
Box plots of tree statistics on transmission trees (*red*) and reconstructed virus genealogies. (*light blue*). Tree height (A), number of cherries per taxa (B), mean internal/external branch lengths ratio (C), Colless Index (D), Sackin’s Index (E), mean branch length (F). The boxes correspond to the first and third quartiles. The upper/lower whisker extends from the third/first quartile to the highest/lowest value which is within 1.5 IQR from the box, where IQR is the inter-quartile range.

Tree height becomes much less informative of network type the bigger the difference is between acute and chronic stage infectivity (Figure 4A). This is because higher acute stage infectivity causes more infections in the acute phase and consequently the epidemic spread is faster. Similarly, mean internal over external branch lengths, which was an important index when constant infectivity was assumed, is less informative if we assume high acute/chronic phase infectivity ratios. Differences observed between ER and BA assuming a constant infectivity profile diminish (Figure 4D). Hence, branch length and tree height measures are less informative of network type when differences in acute-chronic infectivity are considered.

Topological tree measures, i.e., cherries per taxa, and Sackin’s Index, were less affected by differences in acute-chronic infectivities (Figure 4B–Figure 4C). Both these measures, calculated on the possible virus genealogies, still informed about the underlying contact network structure that HIV spread upon.

**Figure 4:**
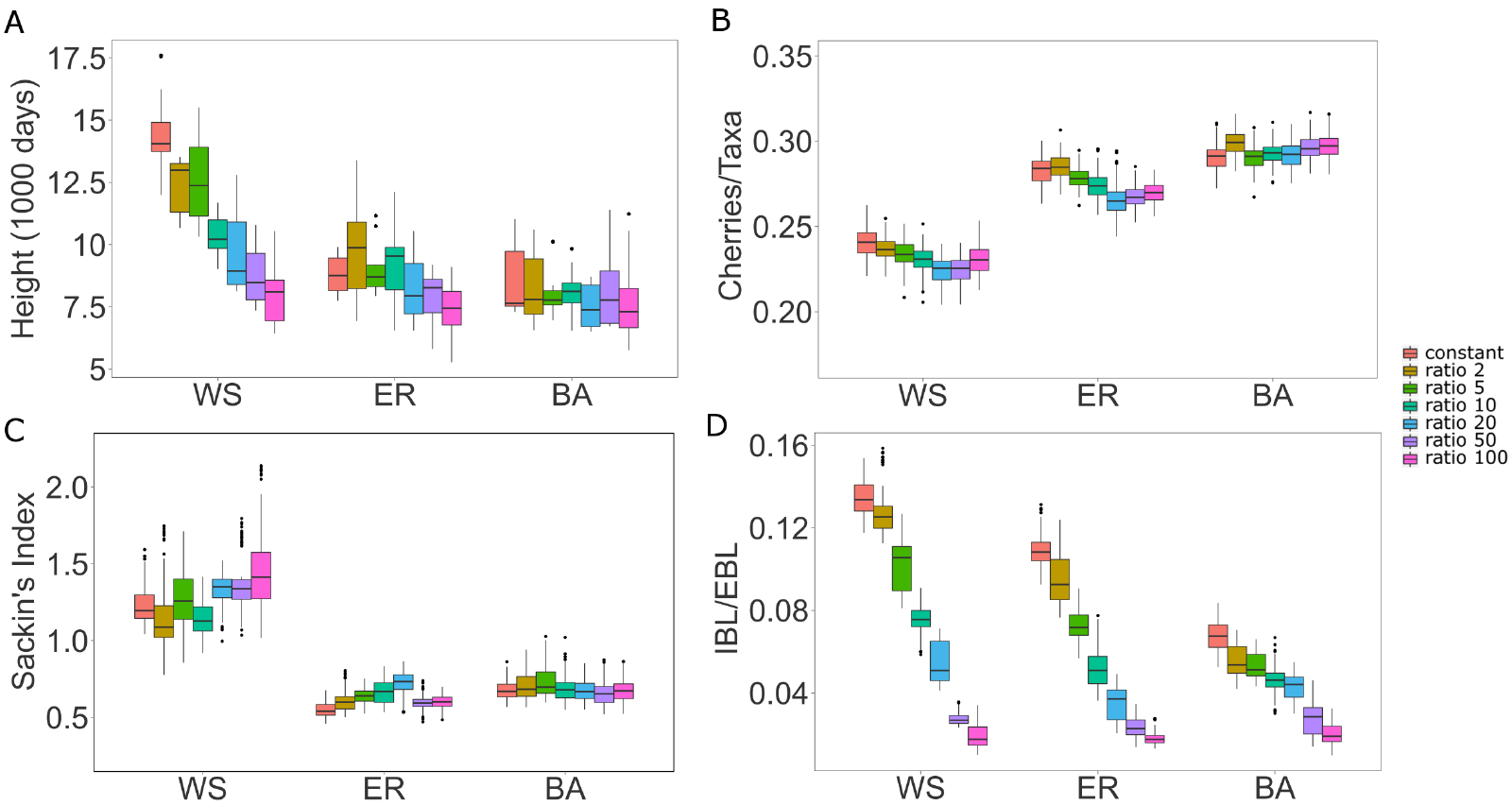
Box plots of tree statistics on virus genealogies under varying infectivity profiles. Tree height (A), number of cherries per taxa (B), Sackin’s index (C), mean internal/external branch lengths ratio (D). Box plots limits are as in Figure 3.

### Individual variability improves network inference

The next stage of introducing realistic host evolutionary dynamics is to model patient specific differences. We did that by introducing variability in the overall infectivity level while keeping the acute-chronic ratio at 10:1 (sigma 0, 3, 10).

Interestingly, while introducing a non-constant infectivity profile diminished genealogical differences between underlying contact network structures, introducing patient variability recovered some of the discriminatory power (Figure 5). While tree height remained with no power to discriminate between networks, internal to external branch length ratios became more discriminative (BA had lowest ratio, ER intermediary, and WS high). Furthermore, both cherries per taxa and Sackin’s Index improved their power to discriminate between contact network structures, and Sackin’s Index could differentiate WS from BA and ER networks.

**Figure 5:**
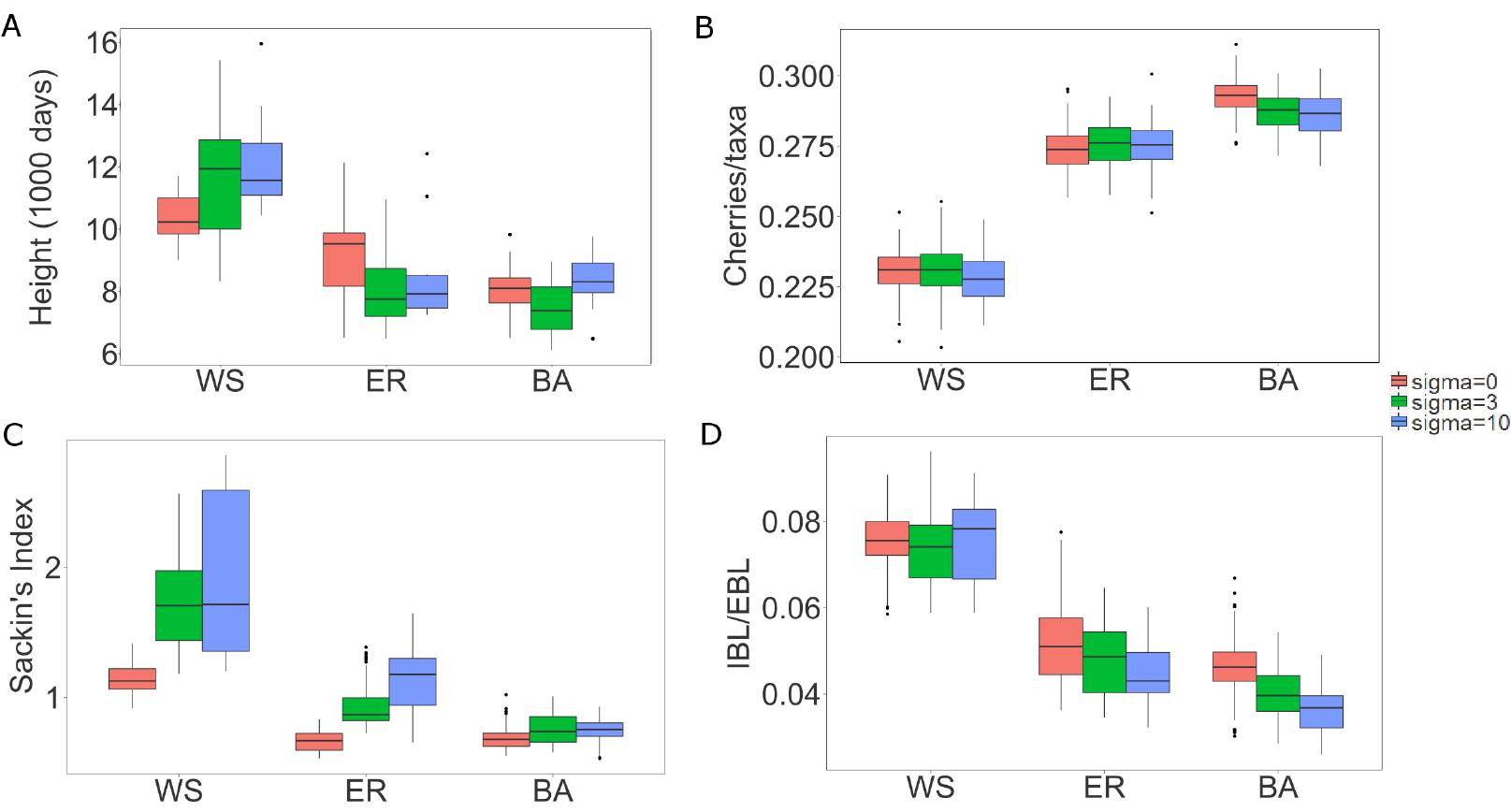
Box plots of tree statistics on virus genealogies for different assumptions on individual heterogeneity. Tree height (A), number of cherries per taxa (B), Sackin’s index (C), mean internal/external branch lengths ratio (D). Box plots limits are as in Figure 3.

### Tree statistics change as epidemics develop

If no influx of susceptibles occurs, the mean branch length increases as trees grow taller because it takes longer time to find uninfected hosts later in an epidemic (Figure 6A). Correspondingly, at 100% sampling of infecteds at any time during an epidemic, the mean branch length increases as a function of total number of sampled infecteds (number of taxa). BA typically produces shorter tree branches than ER and WS as more individuals are sampled (Figure 6B). Thus, if it was possible to sample everyone at time of infection, then the trend of adding longer tips towards the end of the epidemic becomes more pronounced (Figure 6C). The internal to external branch length ratios typically decrease as the epidemic progresses (Figure 6D). This is explained by the fact that branches added later in the epidemic, resulting from chronic donors, divide already existing brandies into shorter segments, BA trees show lower ratios than ER and WS throughout the epidemic, but WS and ER are less distinguishable during an epidemic.

On the topological level, the Sackin’s Index typically decreases as an epidemic matures (Figure 7A), At the end of an epidemic (Figure 3). BA and ER show more unbalanced trees throughout an epidemic and the most imbalanced trees come from WS networks (Figure 7).

Simulations on networks of size 5000 show similar results, for comparison see Figure 6 with Figure S3 and Figure 7 with Figure S4.

Thus, while these statistics are indicative of the underlying contact network, they are confounded by epidemic stage and the size of the susceptible population. Consequently, to be able to infer the underlying contact network from genealogies we must also know what stage an epidemic has reached and the number of susceptibles.

**Figure 6:**
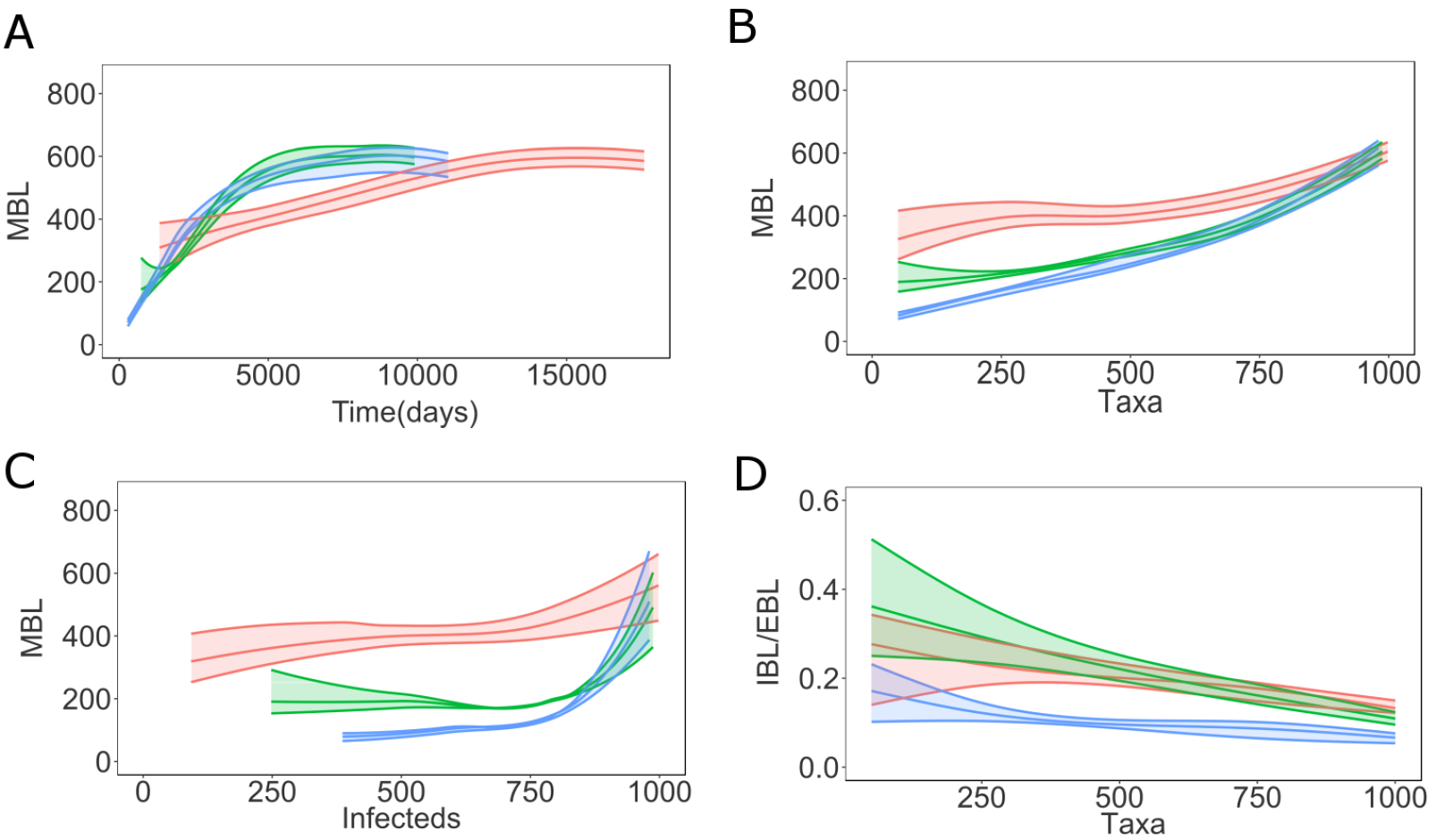
Distance based tree statistics on virus genealogies as epidemic progresses on a network of size 1000. Mean branch length (MBL) as function of tree height (A) and number of infected individuals (B) for simulated outbreaks on networks of size 1000 as epidemics progress, MBL (C) and internal/external branch length ratio (D) as function of the number of taxa for simulated outbreaks on networks of size 1000, The envelopes represent 95% confidence intervals around the medians. The curves are obtained using local regression (LOESS), WS (*red*), ER (*green*), BA (*blue*).

**Figure 7:**
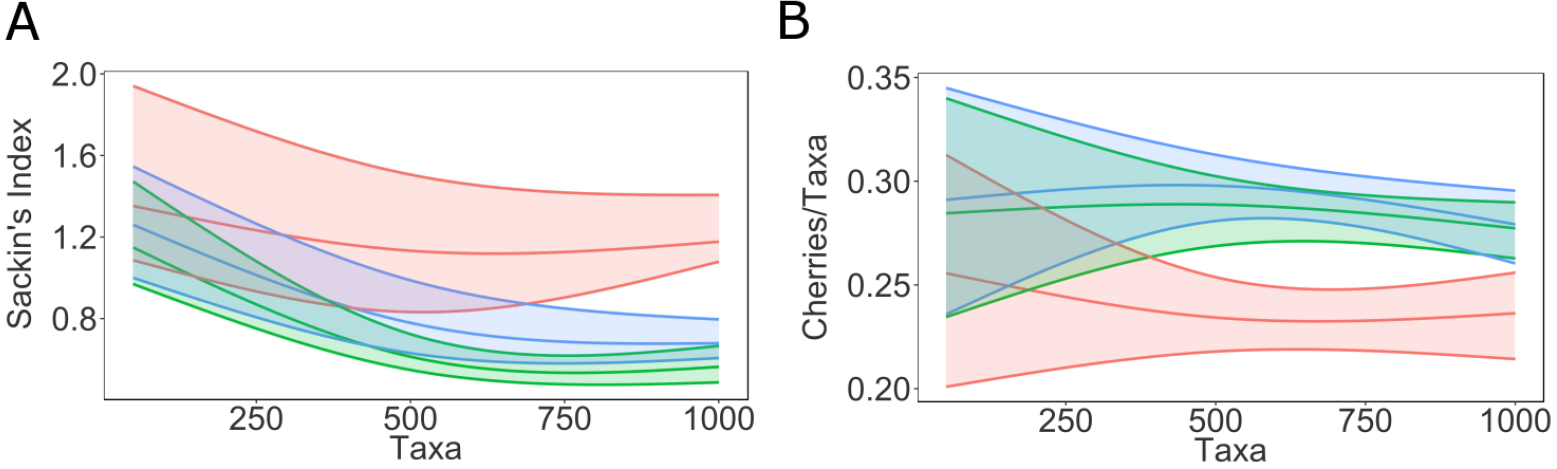
Topological tree statistics on virus genealogies as epidemic progresses on a network of size 1000. Sackin’s index (A) and number of cherries per taxa (B) as function of the number of taxa in networks of size 1000, The envelopes represent 95% confidence intervals around the medians, The curves are obtained using local regression (LOESS), WS (*red*), ER (*green*), BA (*blue*).

### Sample fraction affects tree statistics

While a genealogical tree grows as an epidemic matures, the sampling fraction has no real effect on mean branch length, albeit with smaller sample fractions the estimation becomes somewhat more uncertain due to stochastic effects (Figure 8), Interestingly, lower sampling fraction increases mean branch lengths derived from any underlying contact network (Figure 8), This happens because the remaining branches in the genealogy represent increased numbers of infected hosts. However, this effect does not cause additional confusion over that caused by epidemic stage, as the differences between BA, ER, and WS networks are distinct at all epidemic stages and number of infected, On the other hand, we do not usually know at what stage an epidemic is (i.e., number of actually infected) but only the number of sampled hosts. The mean branch length as a function of number of taxa (Figure 9) could mislead the inference of underlying contact network, especially for small sample fractions. In fact, any branch length or tree height index would be affected by mistaking number of sampled hosts with stage of the epidemic because the number of infected grows faster than the number of sampled early in an epidemic.

The topological indices were also affected by sampling fraction. While general trends (Figure 3) remain constant through the accumulative number of samples over an epidemic, it is again important to know at what stage an epidemic is at time of sampling. Similar to branch length indices, topological indices can be misleading if sampling faction and stage of the epidemic are unknown.

**Figure 8:**
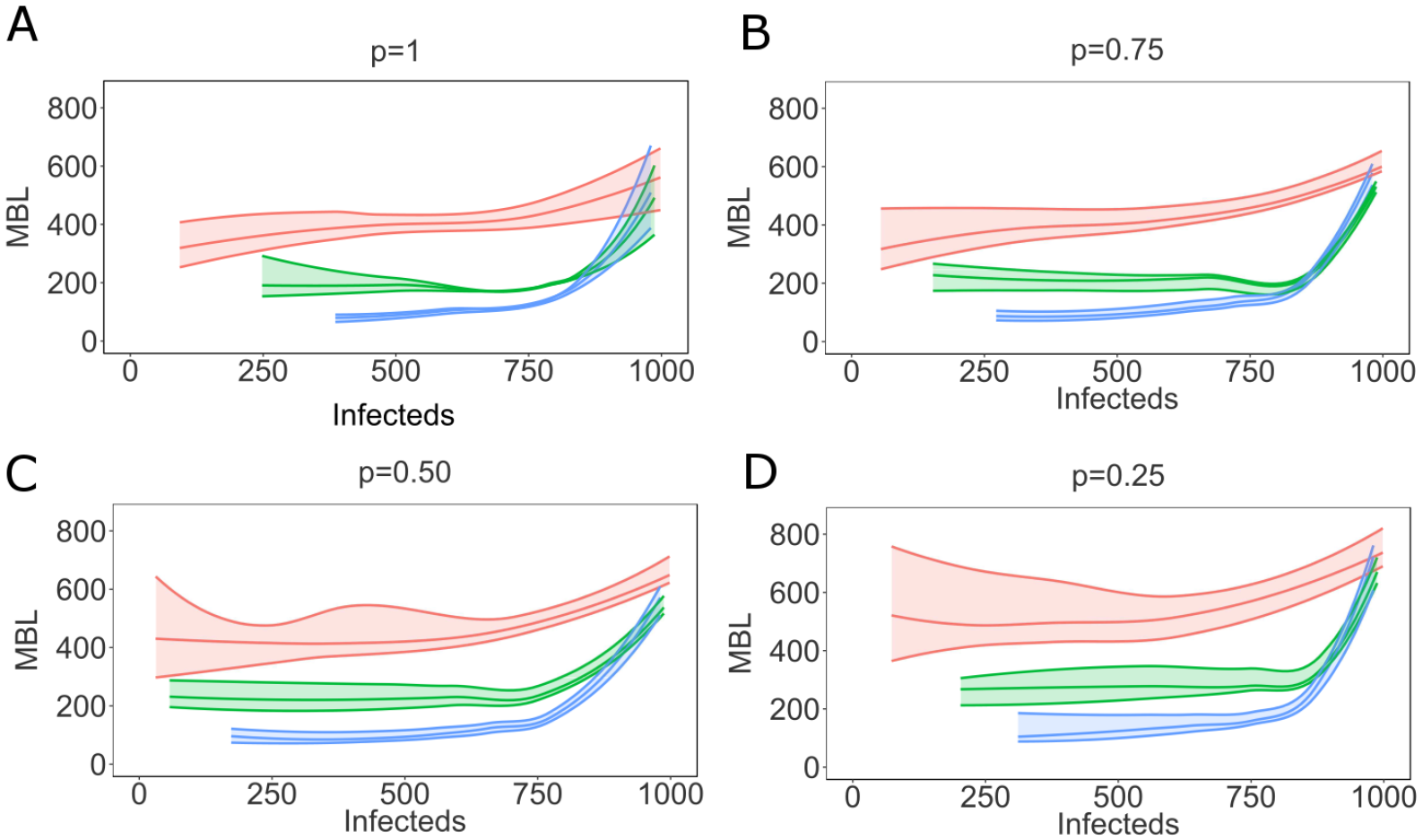
Mean branch length as epidemic progresses varying sampling fraction. The mean branch length as function of number of infectecls, with varying sampling fraction (p=1-0.25), The envelopes represent 95% confidence intervals around the medians. The curves are obtained using local regression (LOESS), WS (*red*), ER (*green*), BA (*blue*).

**Figure 9:**
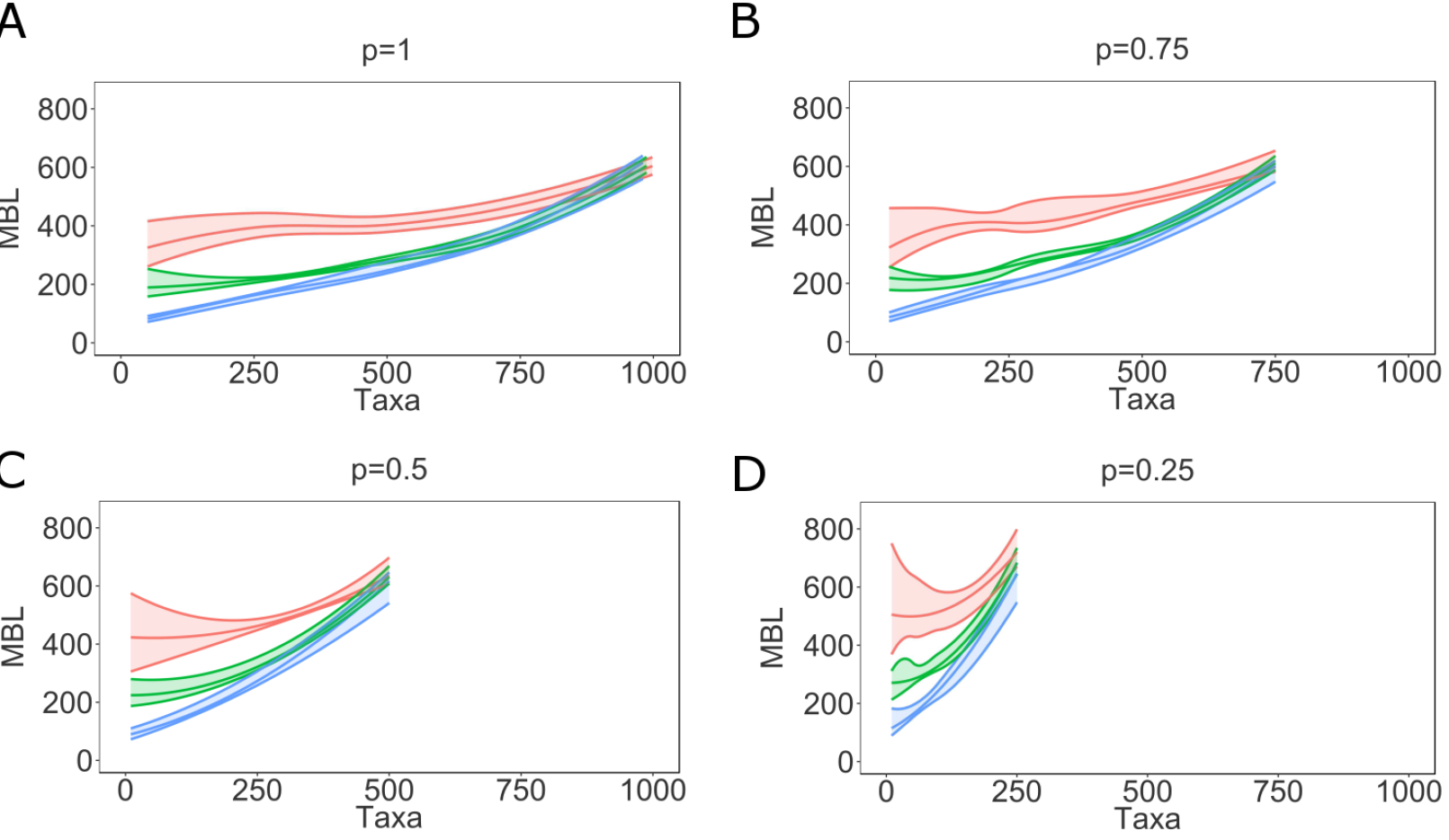
Mean branch length as function of the number of taxa varying sampling fraction. Mean branch length as function of sampled hosts, with varying sampling fraction (p=1-0,25), The envelopes represent 95% confidence intervals around the medians. The curves are obtained using local regression (LOESS), Note that for smaller sampling fractions the envelopes include fewer taxa, WS (*red*), ER (*green*), BA (*blue*).

### ABC inference of transmission network type

We illustrate the performance of the ABC inference on 100 simulated viral genealogies for each network type of size 1000, The parameters were chosen so that the mean degree of each network type was 8, the diagnosis rate was 2.8 years^−1^ (derived from the average time from seroconversion to diagnosis in Sweden estimated in [46]), and infection rate in the acute phase was *λ*_1_ = 0.005. The sampling probability *p* was set at 0.5 because the data from the general HIV Swedish epidemic have coverage of around 50% [46–48].

To investigate model selection performance of the ABC algorithm, we record the number of times that the true model has the highest posterior model probability *P(M*|D) among the three models for the 100 simulated datasets. The algorithm was able to discriminate among the network models quite well. For the first network model (EE), in 78 out of the 100 simulated datasets, the true model had the highest posterior probability among the 3 different network types. For the second model (BA), similar results were obtained; 76 out of the 100 simulated datasets identified BA, Outbreaks on the WS network were miselassified only 1 time out of 100, The corresponding network parameters were estimated reasonably well in most cases (Table 1), An example of obtained parameter estimates is shown in Table 1, where we report mean degree, diagnosis and infection rate for an outbreak on an EE network. The estimation of the removal (i.e. diagnosis) rate was sometimes skewed towards the upper bound, which probably is due to branch elongation induced by the within-host evolution model.

**Table 1:** Parameter estimation for one epidemic spread on an ER network.

| Parameter | Median | 95%CI | True value |
| --- | --- | --- | --- |
| Network parameter, $Np$ | 8.5 | (7.8,8.7) | 8 |
| Removal rate, $\gamma$ | 0.25 | (0.19,0.37) | 0.35 |
| Acute stage infection rate, $\lambda_1$ | 0.008 | (0.002,0.01) | 0.005 |

### Application to data from real epidemics

Inference of epidemic parameters as well as network type becomes more complex in real outbreaks. We consider two genealogies from separate IDU-associated HIV-1 CEF01 and subtype B epidemics in Sweden, respectively. The CEF01 tree was sampled from a rapid outbreak that was imported from Finland [42] around 2003, which was quiescent until the outbreak started in 2006. The subtype B tree was sampled from the more slowly and typical, spreading IDU epidemic in Sweden [41].

While tree indices were different between the trees from the Swedish HIV-epidemic (Figure 10), and superficially in line with what one might expect comparing an outbreak scenario to a more endemic situation, e.g., mean branch lengths were 279 and 913, and tree height 4176 and 10527, respectively, they cannot be directly compared because these trees represent different stages in the respective epidemic. Furthermore, real data is rarely sampled at 100% of all infected or even diagnosed, so comparisons to our simulated overall network differences are difficult to evaluate. Thus, to evaluate genealogies from real epidemics we must consider epidemic stage and sampling fraction (Figure 6–Figure 9)

**Figure 10:**
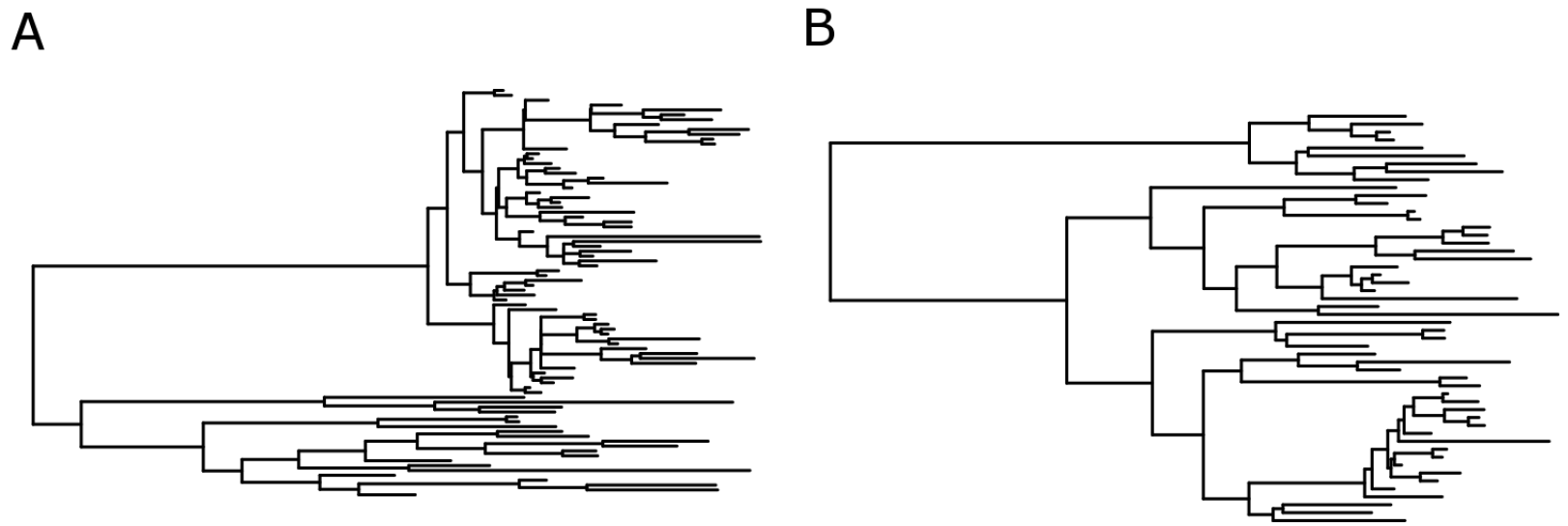
Time-scaled HIV-1 phylogenies from the Swedish epidemic among IDU. A. The genealogy from a rapid CRF01 outbreak, and B the genealogy from a slower spreading subtype B epidemics. Trees were inferred by a Bayesian skyline coalescent model using BEAST 1,8 [43].

In the ABC analysis of the two IDU HIV-1 transmission chains among IDU we have assumed the same epidemiological model as in the simulations (stage-varying infectivity profile with ratio 10:1 acutexhronic, patient infectivity variation (*σ* = 3)) and a sampling fraction of 50%, For the CRF01 IDU outbreak, the susceptible population was assumed to be 200 and in the Swedish subtype B ongoing epidemic it was set to 3000, Results are shown in Table 2.

Overall, convergence was more difficult to achieve in the analysis of the real data and the tolerance levels *ϵ* had to be set to higher values than in the simulation studies. However, the posterior model probabilities seem to indicate that the two outbreaks display different associations to the three network models considered even though there is no single model (among the three network models considered) that can be used to appropriately describe each outbreak.

**Table 2:**
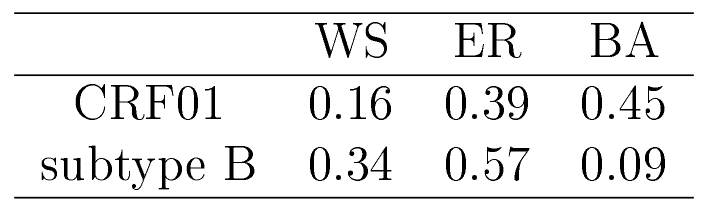
Network type posterior probability for the two Swedish outbreaks.

|  | WS | ER | BA |
| --- | --- | --- | --- |
| CRF01 | 0.16 | 0.39 | 0.45 |
| subtype B | 0.34 | 0.57 | 0.09 |

For the CRF01 outbreak, there is a weak evidence in favor of the BA network type although the EE was also selected with a posterior probability of 39%, with small differences in the parameter estimates. The infection rate in the acute phase was *λ*_1_ = 0.018(0.009, 0.031) vs *λ*_1_ = 0.023(0.0015, 0.036), *γ* = 0.001(0.0006, 0.002) vs *γ* = 0.001(0.0005, 0.002) and the mean degree was 3.2(2.4, 3.6) vs 3.7(2.2, 4.1) in the BA and ER respectively. The HIV-1 subtype B outbreak was mostly (57%) associated with an ER network type. However, there was considerable uncertainty in the parameter estimates *λ*_1_ = 0.0025(0.001, 0.004) *γ* = 0.0003(0.0003, 0.001) degree 1.4(1.3, 2.1).

## Discussion

In this study we addressed several outstanding factors that could affect HIV phylogenetic tree shape in addition to the underlying contact network upon which HIV spreads. While previous studies have evaluated how contact networks affect the resulting tree [10,11,13], they ignored differences between transmission trees and virus phylogenies, varying infectivity over disease progression, among patient infectivity variation, sampling fraction, and epidemic stage. Here, we show that all these factors put further restrictions on what type of phvlogenv one can expect, but also that these additional factors may confound the inference of contact network.

Transmission histories are not perfectly reconstructed by virus phvloge-nies. In fact, it has been previously shown that virus phylogenies have a time bias that elongates external branches, shifts internal nodes backwards, and may cause lineage disordering relative to the transmission history [16], In this study we account for these factors by sampling (many) possible virus genealogies from a transmission history using a recently developed within-host coalescent model [17], Because many virus genealogies may be consistent with one single transmission history, one would expect this factor to add uncertainty to the network inference. However, there is also added signal about transmission times because the within-host diversity changes over the time of infection. Thus, because the degree distributions are different for each network type (Figure 1), transmissions happen after different lengths of infection time, which affects the phylogenetic tree shape. Indeed, we show that the contact network inference from virus genealogies can be quite different than that from transmission trees, and that tree balance differences in fact may be more informative using virus phylogenies. Besides, transmission histories or trees can never be observed, or only partially and then with great uncertainty, which is the main reason for turning to phylogenetic reconstruction in the first place.

It is well known that infectivity is not constant over disease progression, albeit the literature is uncertain about how big the difference is between acute and chronic stage infectivity [45,49], Indeed, we find that varying infectivity affects the expected phvlogenv under different contact networks. In fact, this factor alone seems to diminish phylogenetic differences between contact networks. Somewhat surprisingly, patient variation in infectivity works in the opposite direction, i.e., it amplifies differences in the contact network structures. The result is that virus genealogies do carry a signal of what type of contact network HIV spread upon, but the expectations are different than what one would expect from a naive model where no virus diversity exists and all hosts are described by an identical constant infectivity over their pathogenesis.

We show that any tree index that one would measure is affected by sampling fraction and the stage of the epidemic. We show that phylogenies cannot be meaningfully interpreted without this additional knowledge, as tree statistics otherwise may mislead the inference of contact network. While our results relate to epidemic situations relevant to HIV epidemics, they may also be relevant to other measurably evolving pathogens such as hepatitis C and influenza.

The developed ABC inference framework for network identification and parameter estimation showed discriminatory power and ability to recover epidemiological parameters when applied to simulated data. The model used for validating the ABC algorithm included stage varying infectivity, individual, and within-host variability. For complex models such as epidemic spreads on networks the likelihood function is computationally costly to evaluate and ABC offered a way to perform likelihood-free statistical inference. Furthermore, the use of summary statistics allowed us to study the relationship between readily measurable tree statistics and complex transmission dynamics. The analysis of the two outbreaks from the Swedish HIV epidemic showed that inference on real datasets is typically much harder. As is to be expected, real world networks do not match perfectly with the simplified models considered in this study. In fact, in the ABC algorithm, the proposed parameters values are accepted if the simulated data based on them are close enough to the observed data. If the observed data were generated from a rather different or more complex model, then the simulated data from the candidate model probably will be far away from the observed data. Hence, very few proposed parameter values will be accepted. More realistic models, such as dynamic networks, may be able to better capture the features of the outbreaks, especially those occurring over a long period of time.

## Acknowledgments

This work was supported by the Swedish Research Council (grant number 340-2013-5003) and the National Institutes of Health (NIH) (grant number R01AI087520). We would like to thank Joakim Esbjörnsson for providing useful inputs.

## Appendix SI

## Text S1 Probability to escape infection

We calculated the probability to escape infection, n, as the exponential of the total infectious pressure. Let *X* denote the time to diagnosis (or death from AIDS if never diagnosed) and assume that *X* follows an exponential distribution with rate parameter *γ* i-e. *X* ~ *Exp*(*γ*) In the first model specification (constant infeetivitv) *π* can be written as 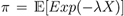 where *λ* is the constant transmission rate. Therefore, 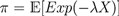=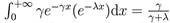.

In the second model specification, which included stage-varying infectivity, let *t_1_* denote the deterministic time spent in the first stage (acute phase), *T_2_* the random time spent in the second stage (chronic phase), represented by an exponential random variable with rate parameter *β*, i.e. *T_2_* ~ *Exp*(*β*), and *t_3_* the time spent in the pre-AIDS stage. Here, the transmission rates in each of the three infection stages are *λ_1_, *λ*_2_* and *λ_3_*, respectively. Thus, the infectious pressure has a different expression depending on when the diagnosis occurs (in the acute, chronic or pre-AIDS stage). The probability *π* is the mean of the infectious pressure calculated over *X* and *T_2_*, We have:

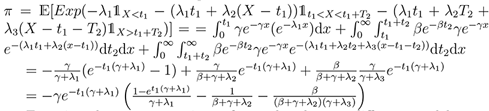

Equating the two expressions of *π* under the two different model specifications, we can calculate the three stage dependent transmission rates *λ_1_, λ_2_* and *λ_3_*, corresponding to a given *λ*. For example, let us assume: *t_1_* = 30,*β* = 1/(365 * 8), *γ* = 1 /(2.8 * 365), *λ_1_* = 100*λ_2_*, *λ_3_* = 100*λ_2_*, *λ* = 0.1%. We obtain: *λ_2_* = 0.0121%

**Figure S1.**
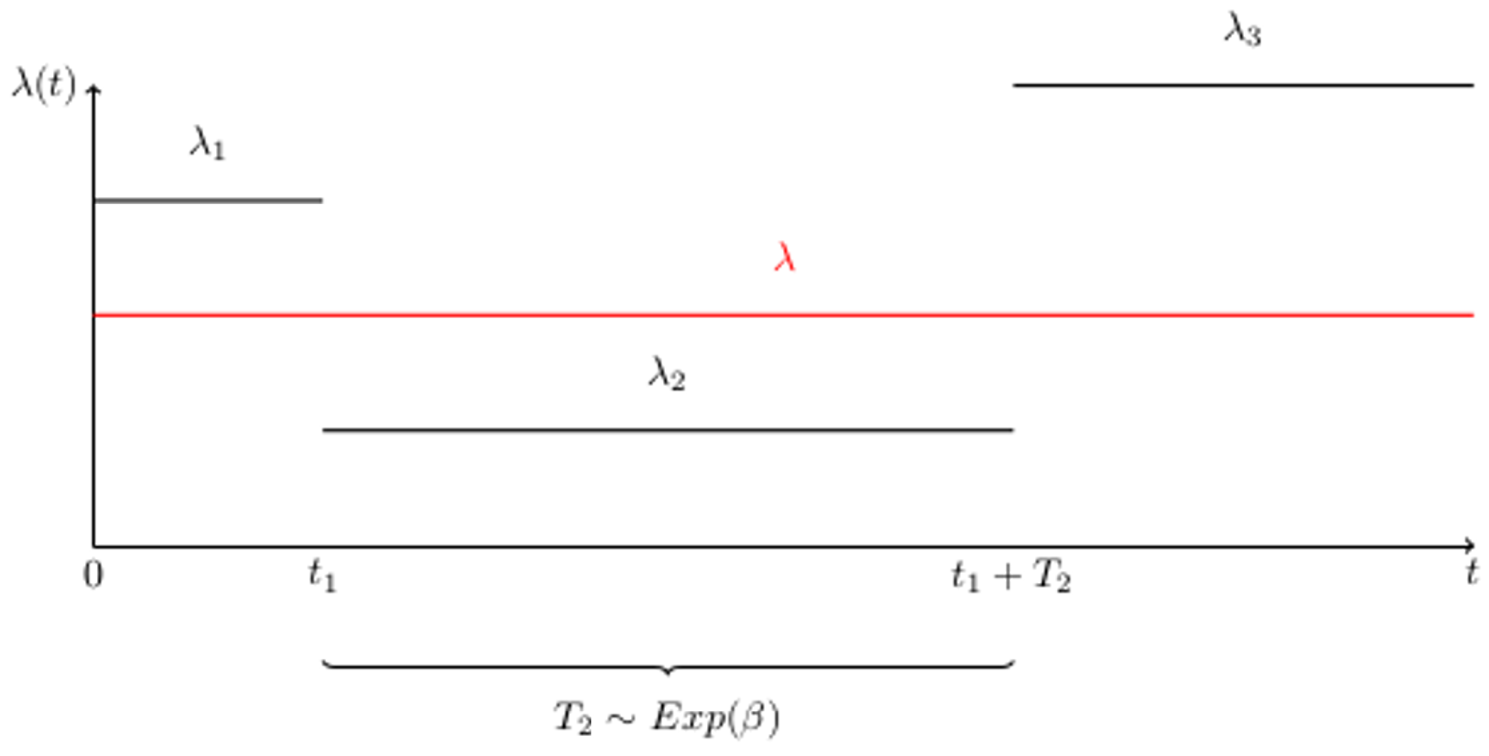
Infectivity profiles. The first two model specifications represented: (i) constant rate (*red line*) and (ii) stage dependent infectivity (*black lines*). The length of the acute phase was assumed constant, i.e. *t_1_* = 30 days while *β* was assumed to be 1/8.

**Figure S2.**
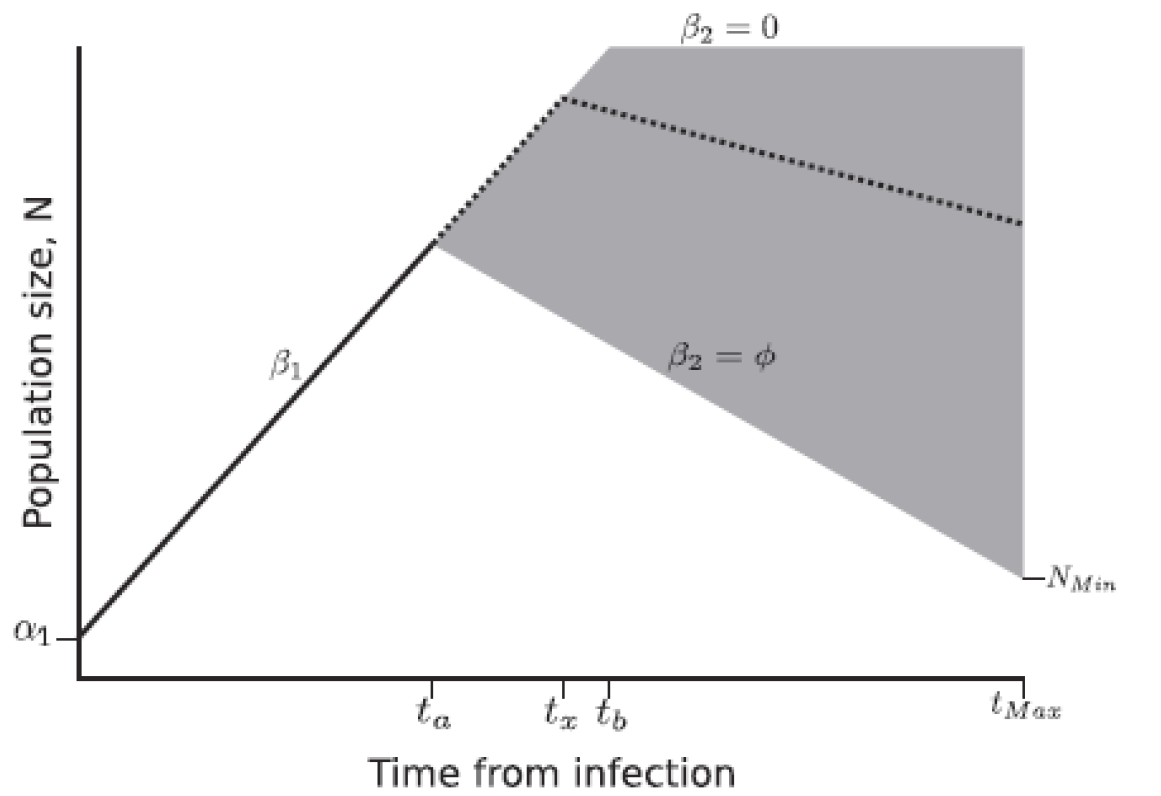
Within-host evolution model. The effective population size is modelled as a two phase linear function: first, it grows at rate *β_1_* until a random peak time *t*_*x*_, after which it decreases or stabilizes. This figure is part of [17].

**Figure S3.**
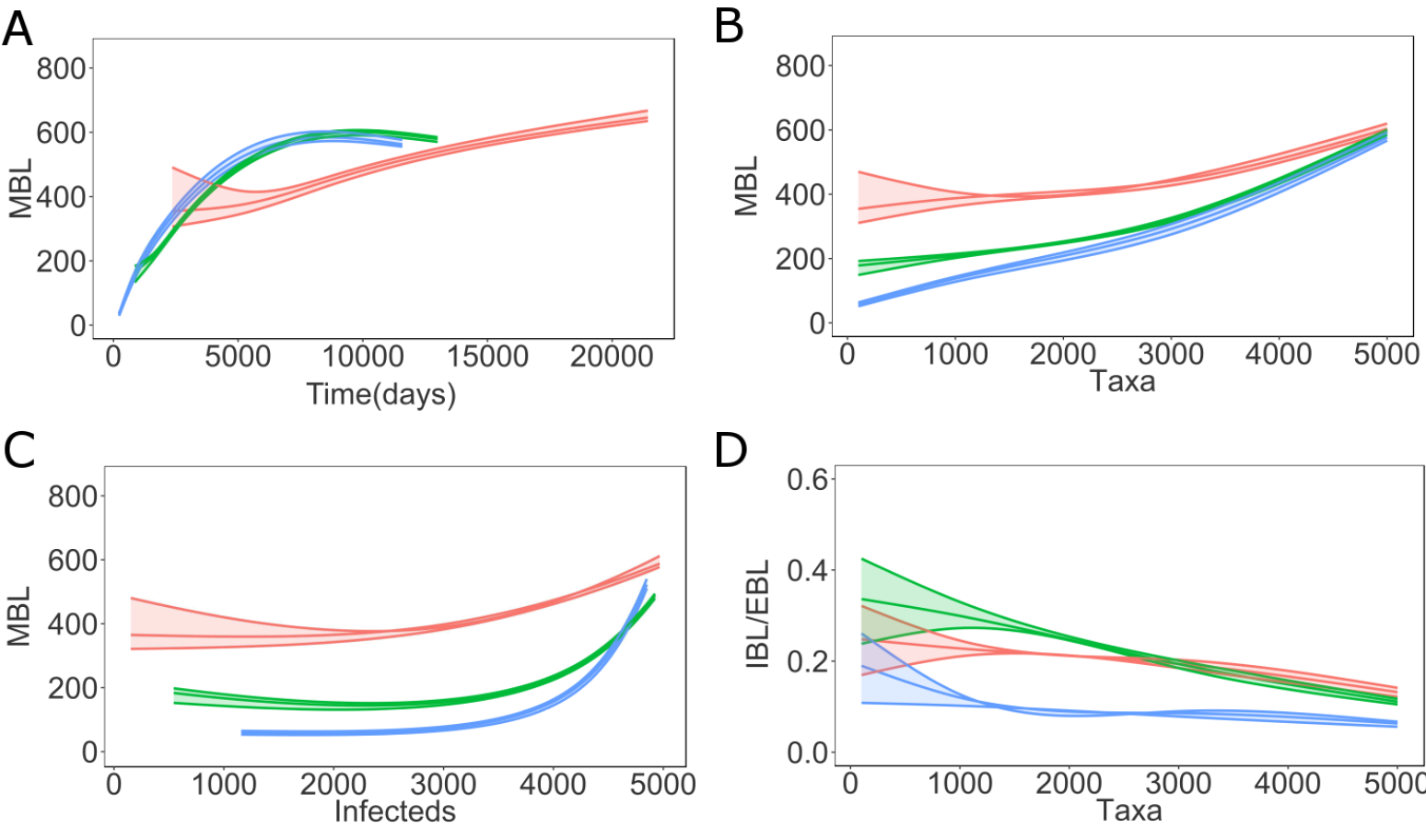
Distance based tree statistics on virus genealogies as epidemic progresses on a network of size 5000. Mean branch length (MBL) as function of tree height (A) and number of infected individuals (B) for simulated outbreaks on networks of size 5000 as epidemics progress, MBL (C) and internal/external branch length ratio (D) as function of the number of taxa for simulated outbreaks on networks of size 5000, The envelopes represent 95% confidence intervals around the medians. The curves are obtained using local regression (LOESS), WS (*red*), ER (*green*), BA (*blue*).

**Figure S4.**
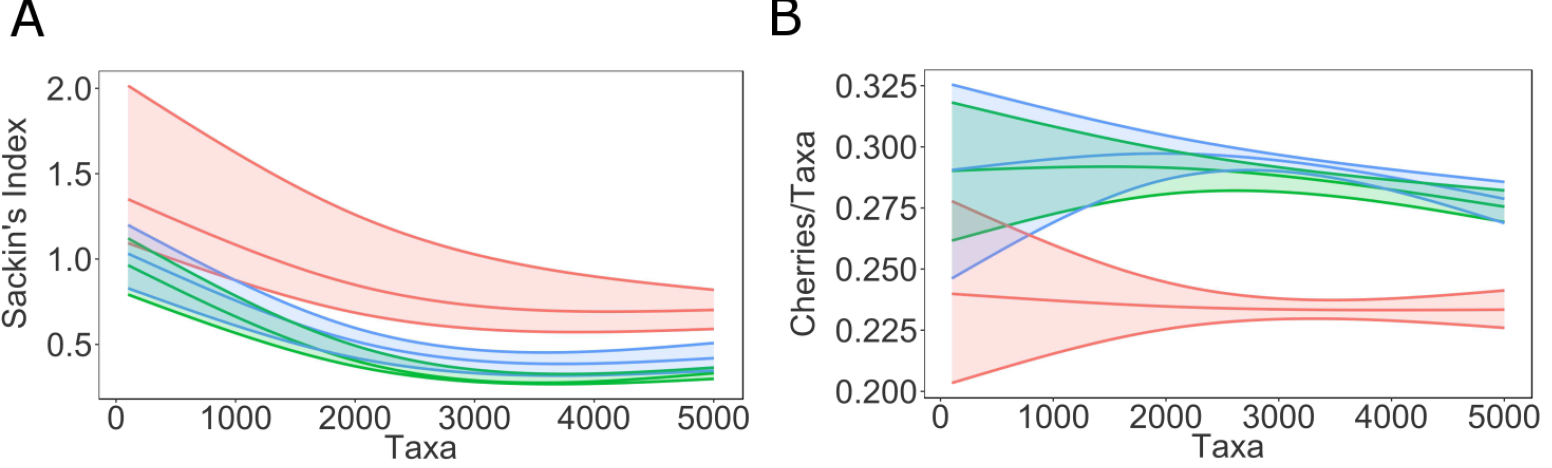
Topological tree statistics on virus genealogies as epidemic progresses on a network of size 5000. Sac.kin’s index (A) and number of cherries per taxa (B) as function of the number of taxa in networks of size 1000, The envelopes represent 95% confidence intervals around the medians. The curves are obtained using local regression (LOESS), WS (*red*), ER (*green*), BA (*blue*).

## Appendix S2

## Text S2 Approximate Bayesian Computation for model choice

Within the AIK ‘-SMC approach [18], particles are first generated from the prior distribution. Particles are then resampled from the obtained sample, and slightly perturbed. From these resampled particles, a new sample is formed, from which again particles are resampled, etc.

The threshold value e for the summary statistic - below which new particles are accepted - is lowered with every newly obtained sample. As a result, the acceptance rate decreases and the acceptance threshold approaches zero with an increase in the number of iterations (resamplings).

Initial *ϵ*-values were estimated as follows. We generated 100 trees and we calculated the summary statistics (indices) and used the standard deviation of this distribution as the initial *ϵ* values. The *ϵ*-values were decreased in an exponential fashion as the sequential ABC scheme progresses

- Initilize *ϵ*
- Set the population indicator *t* = 1
- Set the particle indicator *i* = 1
- If *t* = 1, sample (*m*″, *θ*″) from the prior *π*(*m, θ*) = *π*(*m*)*π*(*θ*|*m*)
- If *t* > 1 sample *m*′ with probabilitv *π*_*t*−l_(*m*′) and perturb *m*″ ∼ *Km*_*t*_(*m*|*m*′) Sample *θ*′ from the previous population {*θ*(*m*″)_*t*−l_} with weights *w*_*t*−l_. Perturb the particle, *θ* ~ *KP*_*t*,*m*″_(*θ|θ*′) where *KP*_*t*,*m*″_ is the particle perturbation kernel. If *π*(*m*″,*θ*″) = 0, repeat this step. Simulate a candidate dataset *x*′ | *f*(*x*~*m*″, θ″) If *ρ*(*x*′,*y*) > *ϵ* repeat this step.
- Set 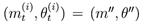 and 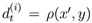), calculate the weight as 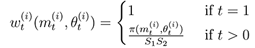

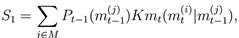

and

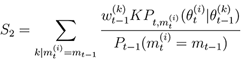
- if *i* < *N* set *i* = *i* + 1 and repeat the previous steps
- Normalize the weights. Obtain the marginal model probabilities given by

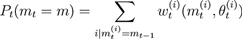

